# Machine Learning-Based Model for Behavioral Analysis in Rodents: Application to the Forced Swim Test

**DOI:** 10.1101/2025.02.18.638880

**Authors:** Andrea Della Valle, Sara De Carlo, Gregorio Sonsini, Sebastiano Pilati, Andrea Perali, Massimo Ubaldi, Roberto Ciccocioppo

**Affiliations:** School of Pharmacy, Center of Neuroscience, Pharmacology Unit, University of Camerino, Camerino, Italy; School of Pharmacy, Physics Unit, Pharmacology Unit, University of Camerino, Camerino, Italy; School of Science and Technology, Physics Division, University of Camerino, Camerino, Italy; INFN-Sezione di Perugia, Perugia, Italy

**Keywords:** Antidepressants, Stress, Rats, Motor Behaviour, Machine Learning, Data Analysis

## Abstract

The Forced Swim Test (FST) is a widely used preclinical model for assessing antidepressant efficacy, studying stress response, and evaluating depressive-like behaviours in rodents. Over the last 10 years, more than 5,500 scientific articles reporting the use of the FST have been published. Despite its widespread use, the FST behaviours are still manually scored, resulting in a labor-intensive and time-consuming process that is prone to human bias and variability. Despite eliminating some biases, existing automated systems are costly and typically only able to distinguish between immobility and active behaviours. Therefore, they are often unable to accurately differentiate the major subtypes of movement patterns, such as swimming and climbing. To address these limitations, we propose a novel approach based on machine learning (ML) using a three-dimensional residual convolutional neural network (3D RCNN) that processes video pixels directly, capturing the spatiotemporal dynamics of rodent behaviour. Our ML model was validated against manual scoring in rats treated with fluoxetine and desipramine, two antidepressants known to induce distinct behavioural patterns. The ML model successfully differentiated among swimming, climbing, and immobility behaviours, demonstrating its potential as a standardized and unbiased tool for automatized behavioural analysis in the FST. Subsequently, we successfully validated our model by testing its ability to distinguish between drugs that predominantly evoke climbing (i.e., amitriptyline), those that preferentially facilitate swimming (i.e., paroxetine), and those that evoke both in a more balanced manner (i.e., venlafaxine). This approach represents a significant advancement in preclinical research, providing a more accurate and efficient method to analyze forced swimming data in rodents. We anticipate that due to its adaptability, in addition to the FST, the model could be applied to various behavioural tests in laboratory animals.

## Introduction

Depressive disorders include a large range of conditions having in common the presence of specific symptoms, such as sadness, emptiness, irritable mood, and loss of pleasure or interest in activities, accompanied by somatic and cognitive changes that significantly affect the individual’s capacity to function [1]. This group of disorders represents a common and global medical problem: it has been estimated that 3.8% of the world population experience depression (approximately 280 million people), including 5% of adults, and it has been projected that this disease will rank first by 2030 [2, 3]. Due to its incomplete responsiveness to pharmacological treatments and the large number of side effects of existing treatments [4, 5, 6], screening of novel antidepressants is still an important practice in current research. A classical preclinical model for screening antidepressant drugs or evaluating depressive-like behaviours in rodents is the Forced Swim Test (FST), originally developed by Porsolt in the 1970s to assess the anti-depressive properties of drugs [7]. The importance of this test is evident from the growing number of scientific publications utilizing it. For instance, a PubMed search using “Forced Swim Test” as a keyword yielded more than 8,600 hits, with over 5,500 just in the last 10 years (see Fig. 1).

**Figure 1.**
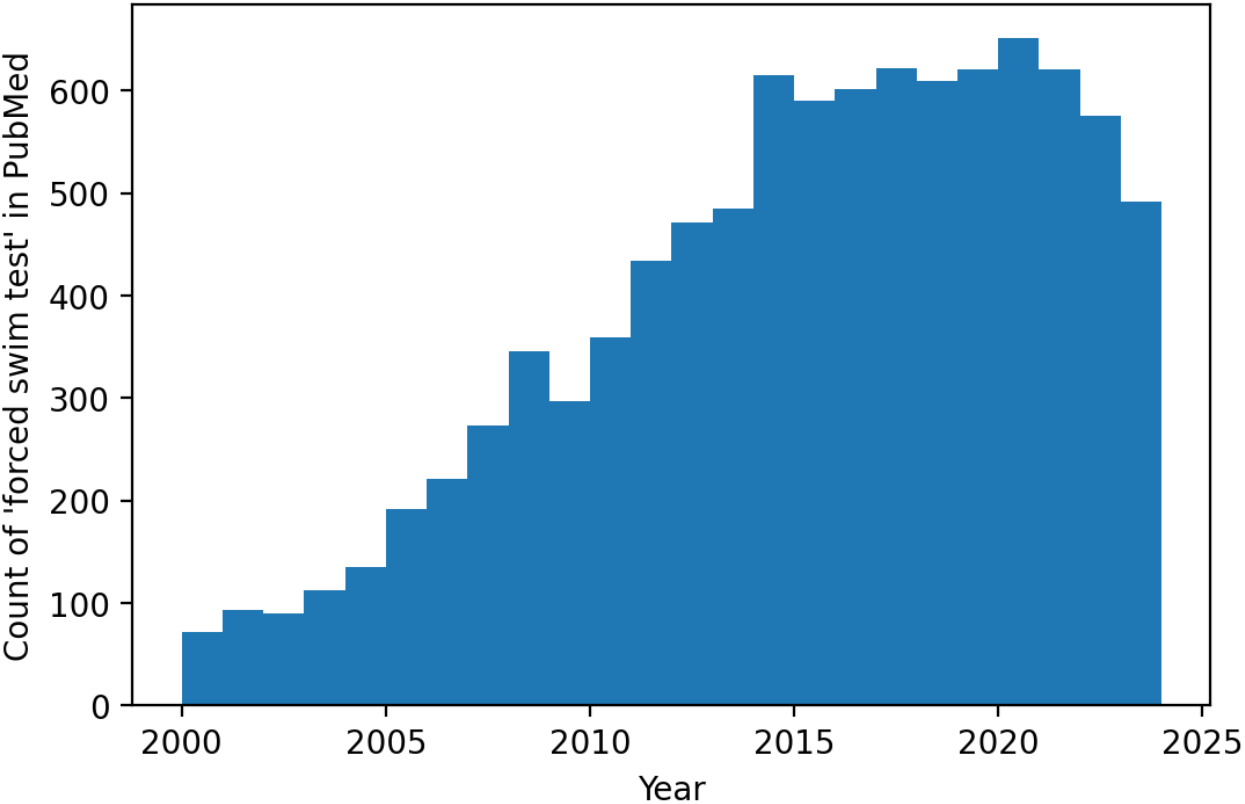
Historical count of the ‘forced swim test’ tag in PubMed. From 2000 to 2024, a PubMed search retrieved more than 8,000 scientific articles using ‘forced swim test’ as a keyword. In particular, 5,500 articles have been published in the past 10 years alone.

The test is based on the observation that rodents immersed in a cylinder filled with water, after initial intense escape-directed behaviour (i.e., swimming and climbing), stop struggling and show passive immobile behaviour (floating with only movements necessary to keep the nose above the water surface), when they learn that escape is impossible. Immobile behaviour is considered a passive coping strategy and is believed to reflect learned helplessness (i.e., behavioural despair) revealing depressive-like behaviour in rodents [8].

Over the years it has been amply demonstrated that the FST is sensitive to all major classes of antidepressant drugs which consistently reduce the amount of immobility time in the test by increasing active escape behaviour [9, 10, 11, 12]. A modified FST originally introduced by Detke and co-workers in 1995 can even distinguish between serotonergic and noradrenergic antidepressants through the distinction of active behaviours into swimming (horizontal movements across the water surface) and climbing (vertical movements against walls) [13, 14, 15, 10]. Lately, the application of this test has expanded to include the evaluation of depressive-like states [16, 17, 18, 19, 20] as well as coping strategies [21, 7, 22] in animal models of psychiatric disorders and stress, becoming one of the most widely used behavioural tests worldwide [22, 23]. Besides having good predictive validity, the FST is straightforward to conduct and requires minimal specialized equipment. Nevertheless, there are still some major hurdles associated with it.

In a large majority of the studies, animals’ behaviours have been evaluated by hand by a trained annotator, and less frequently automated systems based on video analysis have been employed. Both methods have relevant but different limitations. The manual approach is prone to human subjectivity, making it difficult not only to replicate findings across laboratories but also to maintain standardized scoring methods over time in the same laboratory. Moreover, manual scoring requires a large amount of training time, it is very time-consuming, and even scoring by the same researcher shows changes over time due to the experience gained during work or other potential bias factors [24, 25, 26]. On the other hand, the currently available automatic systems are costly and do not allow adequate detection of the specific kind (i.e., swim vs climbing) of behaviour performed by the rodent. Indeed, these methods extrapolate the degree of mobility of the animal by evaluating the frame-to-frame variations and/or the position and velocity of the centroid of the animal or other key points (such as nose and tail) [27, 28, 29, 30, 31, 32, 33].

Considering this general framework, there is a clear need to develop alternative methods to overcome the limitations outlined above and to characterize rodent behaviours more effectively [34]. Having this purpose, here, we propose a new approach based on machine learning (ML) implemented via deep neural networks.

To decrease the human bias given by the selection of some specific extracted features, we work with the information given by the frame, taking into account the dynamical evolution encoded in temporal sequences of frames. Indeed, rodents’ behaviours feature complex sequences of dynamic actions, and therefore the information provided by a single frame is not sufficient to appreciate the spatiotemporal nature of the behaviour and to classify it properly. Hence, in our approach, small portions of the video are preprocessed into 3D-tensors [35]. The ML architecture chosen to process such kind of data is a 3D residual convolutional neural network (3D RCNN). Indeed, these architectures have already proven successful in the literature on human action recognition [36, 37]. For the training and the later testing of the ML model, datasets of video-recorded FST were collected and, when necessary, manually scored in our laboratory.

Once the ML algorithm was trained and validated, we tested it by comparing its outcome with that produced by the human manual scoring on rats under the influence of the selective serotonin reuptake inhibitor fluoxetine and tricyclic antidepressant desipramine. The first is known to predominantly increase swimming while the latter evokes a marked enhancement in climbing behaviour [38, 15, 14, 39]. Once a good performance of the ML algorithm was demonstrated, a second FST experiment with different antidepressants was carried out to confirm the ability of our ML algorithm to properly identify the antidepressant based on the behavioural repertoire evoked and described in the literature [14, 40, 41, 42]. Results demonstrated that our ML algorithm successfully discriminates among the three behaviours, proving to be a standardized, unbiased, and objective method for behavioural analysis of the FST in rats. If appropriately trained, we note that this model can be applied to analyze various behavioural tests including but not limited to the open field, fear conditioning, and drug-induced withdrawal symptoms.

## Results

To develop the ML model, we used an in-lab-made training dataset consisting of 78 video-recorded forced swimming tests. To mitigate the disadvantages associated with running the study in different environmental settings, here we have provided geometrical details on how to perform the experiment (Table 1).

**Table 1.** Principal parameters used for the experiments. Relative distances between cylinder and camera to record the videos are reported.

|  | **Distance [cm]** |
| --- | --- |
| Camera – floor | 25 |
| Camera – back cylinder | 120 |

Those videos were divided into 10,062 sub-videos of 3 seconds and mapped into a standardized 3D tensor through a phyton script written using OpenCV module [43]. For each sub-video, the main behaviour was identified by two experienced researchers who labeled the behaviours as immobility, swimming, or climbing. An example of the 3D tensors for each class of behaviour is shown in Fig. 2.

**Figure 2.**
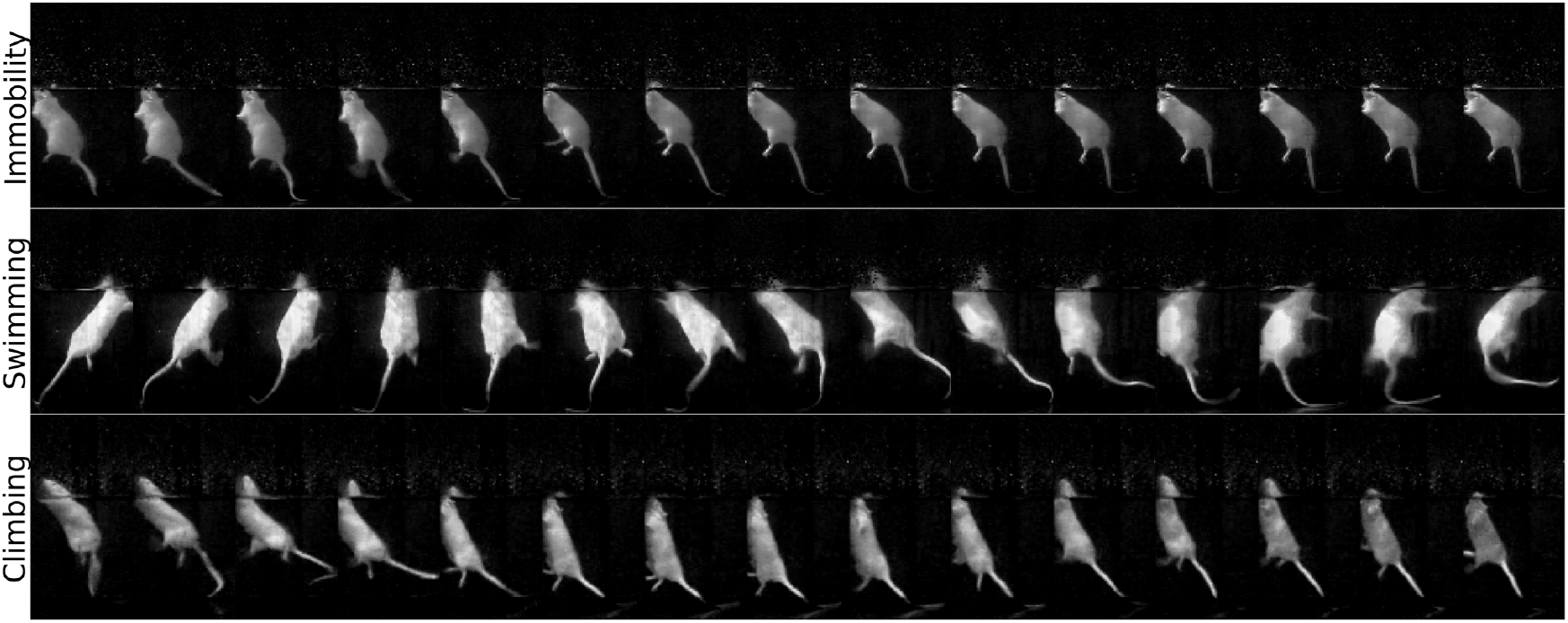
Example of 15 frames extracted from the 3D-tensor, for each behaviour. Each 3D-tensor contains 75 frames. For the present representation, one frame was shown every five frames.

Then, this training dataset was split into 90% train and 10% validation sets, for the training of the ML model.

The ML model trained is a 3D RCNN permitting the extraction of complex spatio-temporal features useful for the classification task. The model was coded using the Keras library [44] with tensorflow backend [45]. The architecture is sketched in Fig. 3.

**Figure 3.**
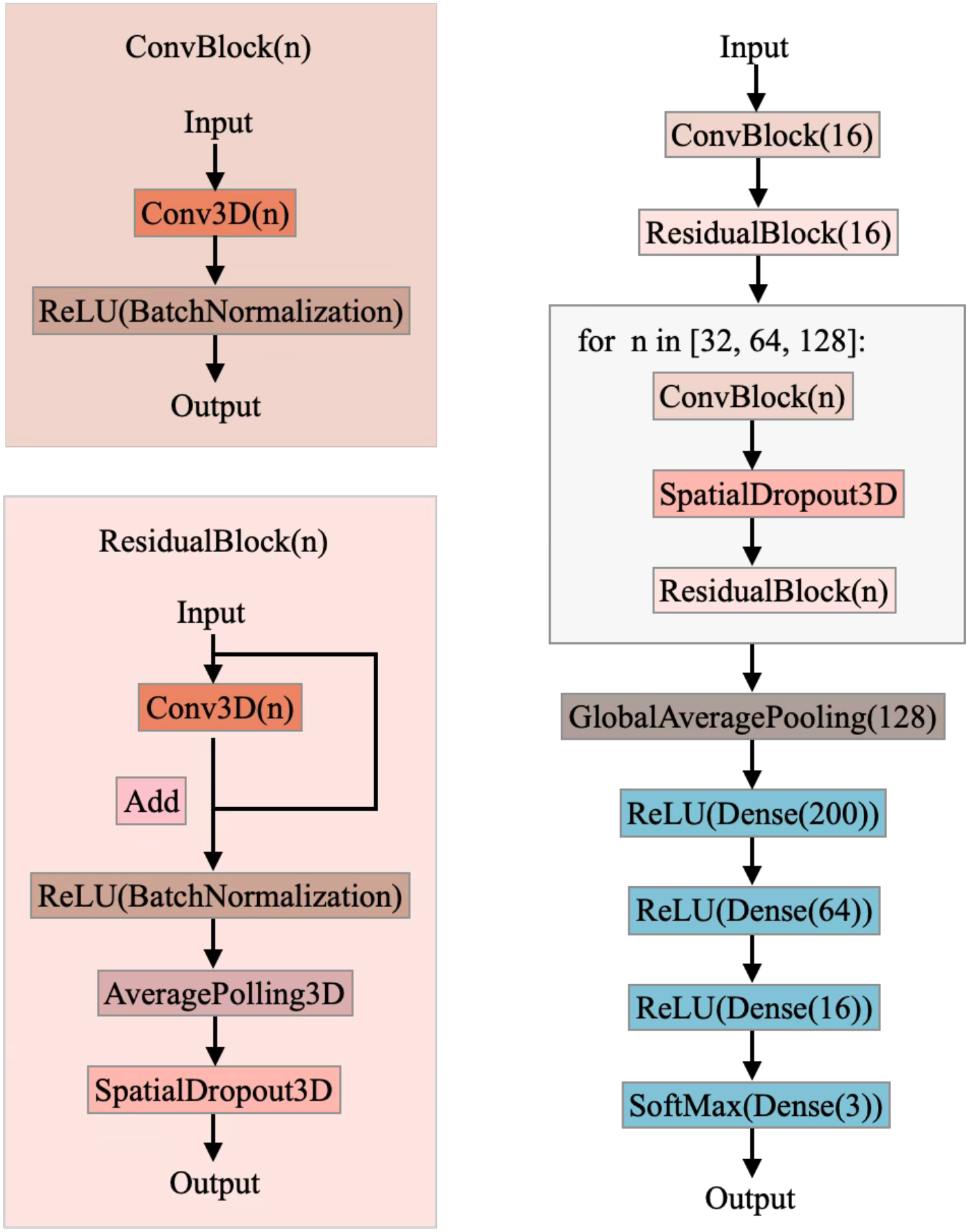
The model architecture used for behaviour classification. The model is composed of a cascade of 3D convolutions organized in residual blocks to extract the spatio-temporal features of the data. Then, after an average global pooling, a series of dense layers was used to learn non-linear combination of the extracted features. Finally, a three-node softmax layer is used to produce a probability distribution of the possible behaviours label. The behaviour corresponding to the highest probability is the output of the model. We set the kernel size of the 3D convolutions to (3,3,3), the kernel size of the 3D average pooling to (2,2,2), and the dropout value to 0.1. The variable n in the convolutional neural layer is the number of its output channel.

The model was trained by using Adam optimizer [46] on 120 epochs, and we used early stopping on validation accuracy as regularization (accuracy of 88.89% on validation set, Fig. 4A).

**Figure 4.**
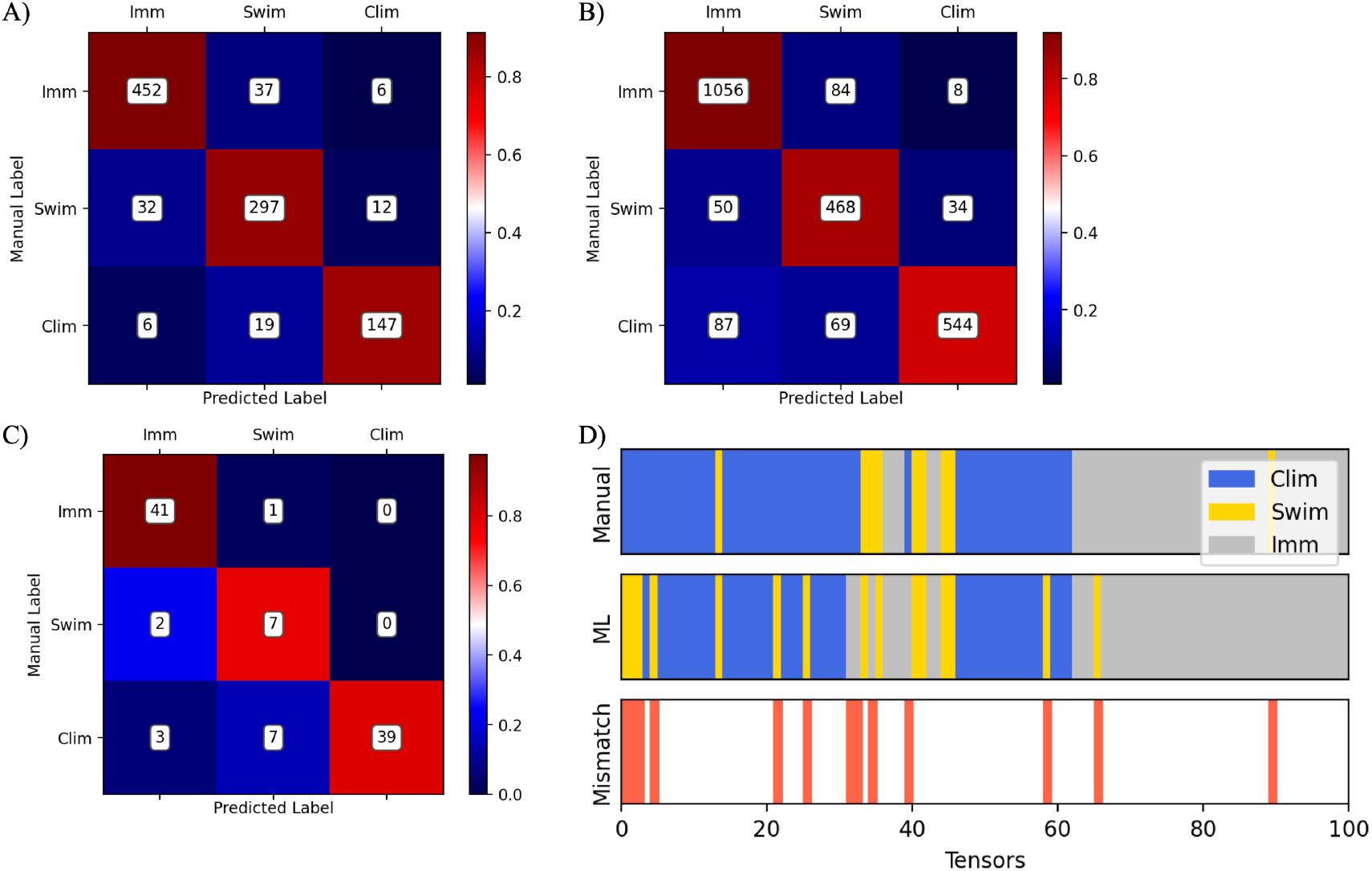
Performance of the ML scoring compared with manual scoring. The confusion matrix on the validation set (**A**). The confusion matrix on the sum of the drug experiment videos (**B**). As an example, a video has been selected to show the concordance between the two scoring methods via the confusion matrix (**C**) and via the time plot in which any mismatch between the two methods is also shown (red bars). Each 5-minute video is split into a sequence of 100 3-second tensors, used as inputs for the ML algorithm (**D**).

### Comparison of the ML algorithm with the human manual scoring on the effects of fluoxetine (FLX) and desipramine (DMI) in the FST

The ML labeling and the manual scoring are in good agreement, as noticeable from the confusion matrix of the complete dataset in Fig. 4B, and the 86 ± 4 % accuracy measured on the videos (mean ± standard deviation). Fig. 4C and 4D show an example of a video labeled by the annotator and by the ML: the confusion matrix of the single video and the time-plot of the two behaviour recognitions (by the annotator and the ML) with their differences (red bars), respectively.

There are strong correlations between the two scoring methods (Fig. 5) as confirmed by the following analyses. For the total times classified in each class, as measured by the Pearson’s correlation, the following scores are obtained: immobility: *r* = 0.918; swimming: *r* = 0.858; climbing: *r* = 0.955 (Fig. 5). ANOVA analyses are run on each behavioural data scored using either method in order to compare them and verify whether the ML algorithm could recognize a statistically significant difference in response to antidepressant as human manual scoring would do.

**Figure 5.**
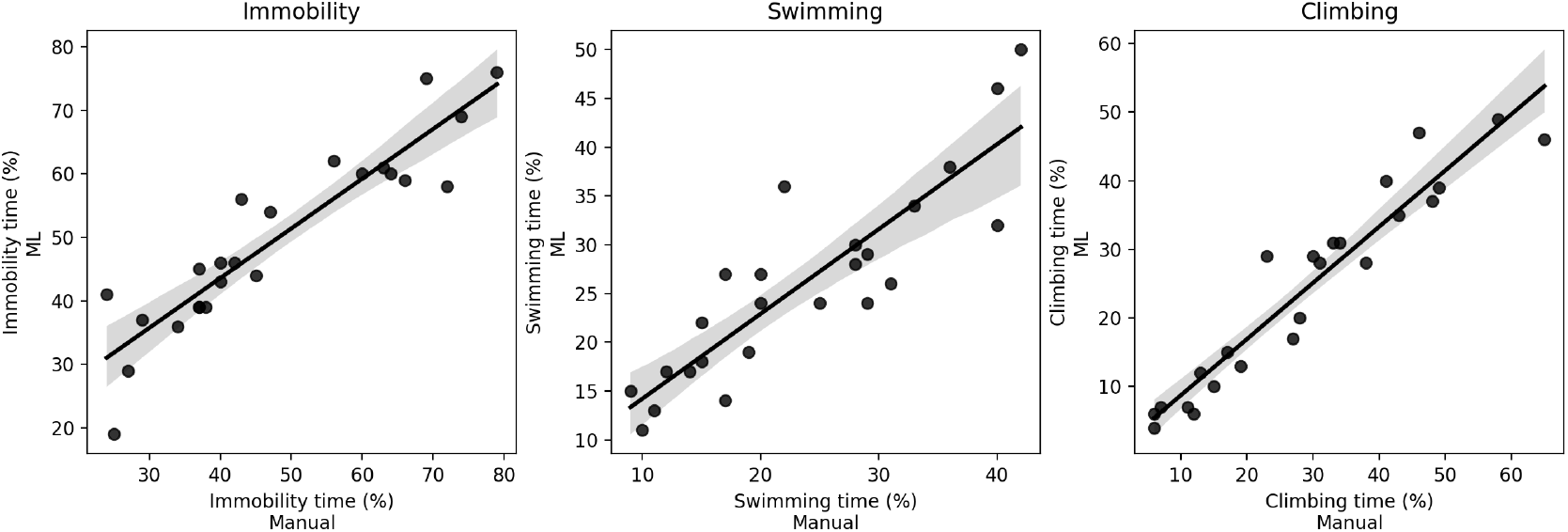
Comparison between the behaviour times of behaviour measured with the two scoring systems. Scatter plots of the percentage time performing the three behaviours predicted by the trained model show the positive correlation between manual and ML labeling systems. As measured by a Pearson’s correlation, the following scores are obtained: immobility r = 0.918 ; swimming r = 0.858 ; climbing r = 0.955.

ANOVA demonstrated no statistically significant difference between the manual and the ML scoring, whereas a statistically significant effect of the drugs in determining the specific type of behaviour expressed by the animals was detected. The time spent immobile significantly decreases in rats treated with DMI and those treated with FLX compared to the control group (Fig. 6A). The analysis also shows that the animals treated with FLX exhibit higher time performing swimming compared to the vehicle-treated animals (Fig. 6B). Whereas the animals treated with DMI show enhanced time performing climbing behaviour compared to the control group (Fig. 6C). Both scoring methods measured these statistically significant differences.

**Figure 6.**
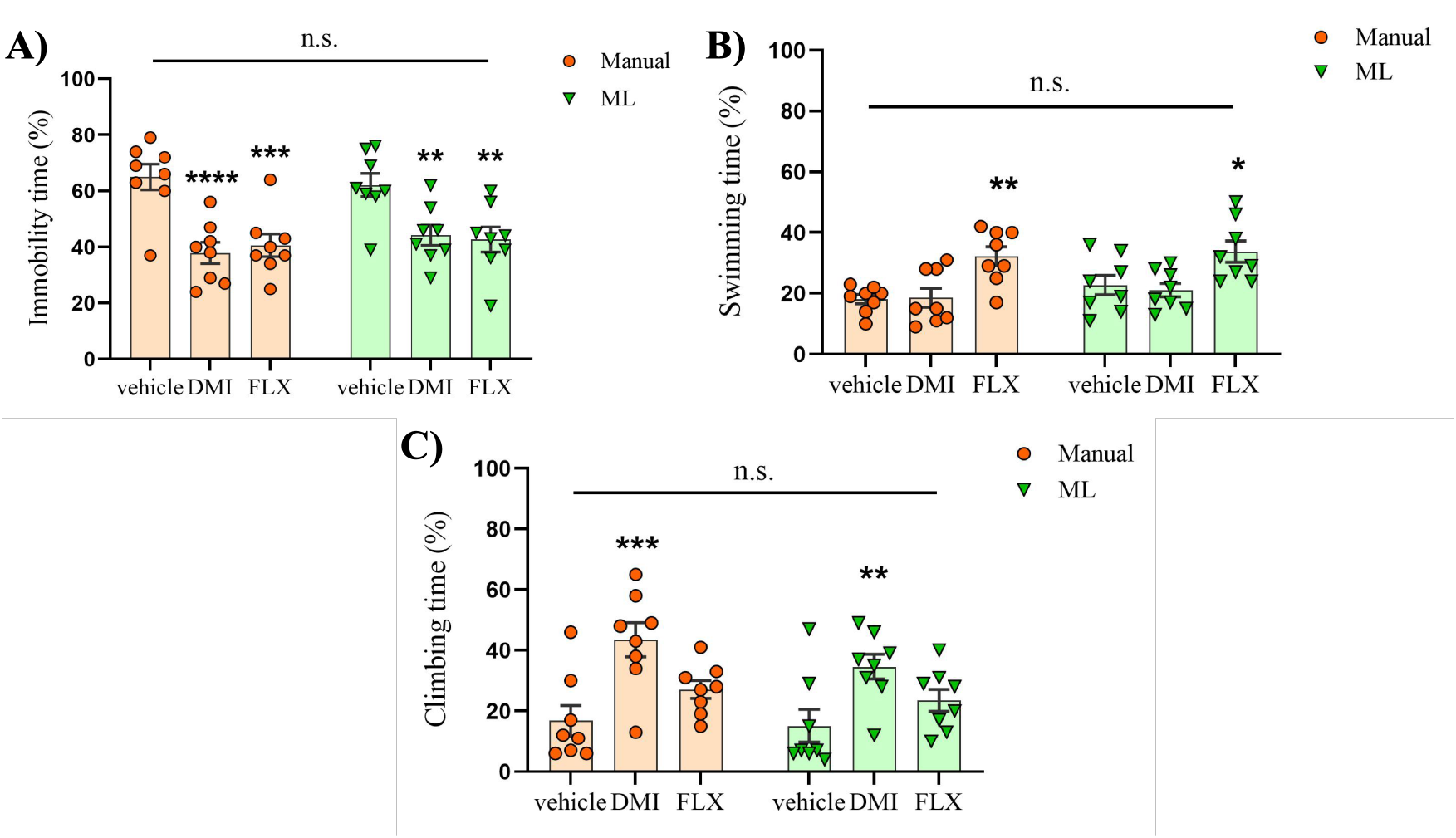
Measurement of the effect of the drugs on FST behaviour measured with ML algorithm and manual scoring. Comparison between the manual and the ML algorithm scoring following drug treatment. When the ML algorithm was compared with manual scoring, no statistically significant differences were detected between the two methods for any behaviour assessed in the FST (Immobility: F(1,42) = 0.31, p = 0.579; Swimming: F(1,42) = 1.5, p = 0.227; Climbing: F(1,42) = 1.64, p = 0.208). Consistent with well-established literature on the effects of FLX (n = 8) and DMI (n = 8) in the FST, both methods successfully identified: (**A**) a decrease in immobility time (F(2,42) = 19.42, p < 0.0001) in rats treated with antidepressants compared to the vehicle group (Manual: vehicle vs. DMI, p < 0.0001; vehicle vs. FLX, p = 0.0003. ML: vehicle vs. DMI, p = 0.007; vehicle vs. FLX, p = 0.003) (**B**) the SSRI FLX preferentially increased swimming behaviour (F(2,42) = 13.34, p < 0.0001. Manual: vehicle vs. FLX, p = 0.002. ML: vehicle vs. FLX, p = 0.018); and (**C**) the TCA DMI preferentially increased climbing behaviour (F(2,42) = 13.02, p < 0.0001. Manual: vehicle vs. DMI, p = 0.0003. ML: vehicle vs. DMI, p = 0.008). Two-way ANOVA followed by Dunnett’s multiple-comparisons test was used. Data are presented as mean ± s.e.m. *p < 0.05, **p < 0.01, ***p < 0.001, ****p < 0.0001 vs. vehicle treatment.

### ML algorithm evaluation of the effects of amitriptyline (AMI), paroxetine (PRX), and venlafaxine (VLX) in the FST

Based on previous results we predicted that our ML is able to discriminate between the various classes of antidepressants. To confirm this assumption, we used our ML algorithm to analyze the behavioural responses to three well-known antidepressants, amitriptyline, paroxetine, and venlafaxine, characterized by distinct mechanisms of action.

The analysis reveals a decrease in the immobility time in the animals treated with AMI and VLX compared to the vehicle group (Fig. 7A). Rats that received PRX and VLX significantly increase their time spent performing swimming (Fig. 7B), while animals treated with AMI significantly perform more climbing compared to the vehicle group (Fig. 7C).

**Figure 7.**
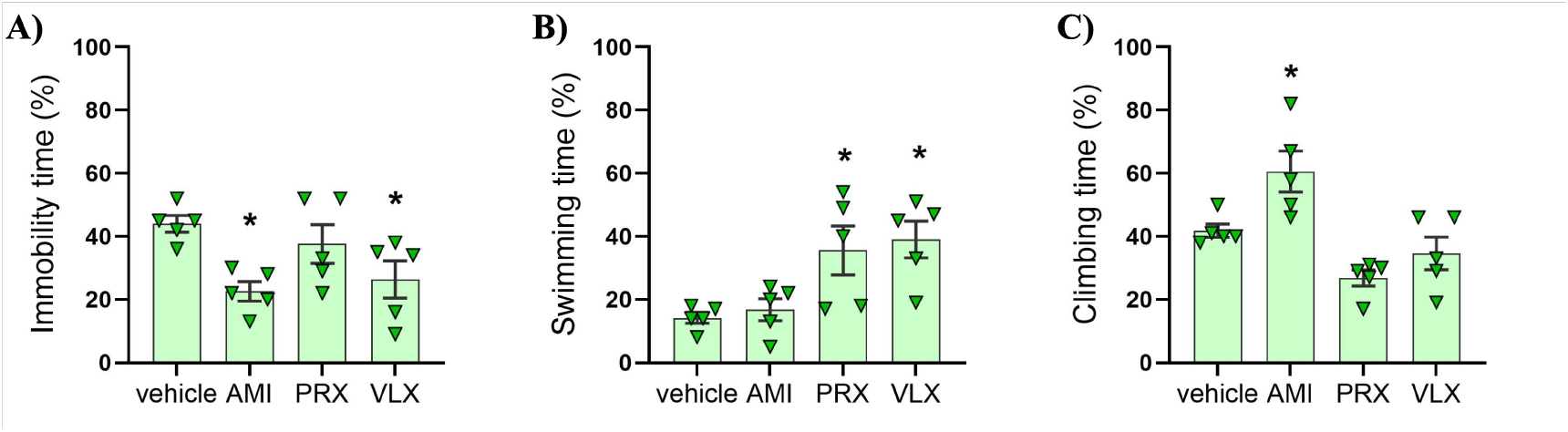
Measurement of the effect of the drugs on FST behaviour measured with ML algorithm. The ML algorithm accurately categorizes the effect of different classes of antidepressants in the FST. According to the literature, our ML model assigned the TCA AMI (n = 5) high levels of climbing, whereas, for the SSRI PRX (n = 5), the model detected increased swimming. In the case of VLX (n = 5), which, compared to TCAs and SSRIs, has a more balanced noradrenergic/serotonergic profile, our model indicated a corresponding balance between swimming/climbing performance. (**A**) Immobility (F(3,16) = 4.48, p = 0.018. vehicle vs. AMI, p = 0.014; vehicle vs. PRX, p = 0.656; vehicle vs. VLX, p = 0.044); (**B**) Swimming (F(3,16) = 5.93, p = 0.006. vehicle vs. AMI, p = 0.97; vehicle vs. PRX, p = 0.027; vehicle vs. VLX, p = 0.011); (**C**) Climbing (F(3,16) = 10.55, p = 0.0005. vehicle vs. AMI, p = 0.023; vehicle vs. PRX, p = 0.075; vehicle vs. VLX, p = 0.541). One-way ANOVA followed by Dunnett’s multiple-comparisons test was used. Data are presented as mean ± s.e.m. *p < 0.005 vs the vehicle treatment.

These results are in line with data from the FST literature demonstrating that PRX and VLX preferentially increase the swimming time [14, 42], while AMI increases the climbing time in the rat [40, 41].

## Discussion

The FST is a well-validated animal test fundamental to understanding the underlying pathophysiology of depressive disorders, assessing stress coping strategies, and evaluating the efficacy of existing and novel antidepressant drugs. However, despite the large use of this test in laboratory animals, a bias-free, accurate, reproducible, and less labor-intensive method to score the FST is still lacking.

In the present paper, we describe the development of an algorithm based on a 3D RCNN ML model, that automatically recognizes immobility, swimming, and climbing, the three main types of behaviour in the FST. Specifically, using a FST video dataset recorded in our laboratory, we first trained the ML algorithm to recognize these behaviours. Afterward, we validated it by applying it to the analysis of two additional datasets from experiments with antidepressants. The first drug experiment was labeled both by hand and by using our ML algorithm, while the second experiment was analyzed only by the ML algorithm.

The results demonstrate that once trained the model can correctly categorize and quantify the three behaviours in high accordance with the manual scoring of the two trained researchers. Moreover, data obtained following drug treatments demonstrated that the model is able to discriminate between different classes of antidepressants. In fact, in accordance with the literature the model assigned high levels of climbing to desipramine and amitriptyline, two predominantly noradrenergic tricyclic antidepressants (TCAs). Whereas when fluoxetine and paroxetine, two selective serotonin reuptake inhibitors (SSRIs), were tested the model detected higher levels of swimming than climbing [13, 14, 15, 10]. Venlafaxine, compared to TCAs and SSRIs, has a more balanced noradrenergic/serotonergic profile. Accordingly, our model assigned to this drug a more balanced swimming/climbing performance. Finally, for all the antidepressants tested our ML model identified a significant increase in mobility, which is also consistent with the results of manual scoring reported in the literature [13, 14, 15, 10].

Compared to human manual scoring and the current automatic systems, our 3D RCNN model presents several advantages. For instance, we estimate that the time necessary to label a 5-minute video by hand is at least 30 minutes, while our ML approach, considering also the whole preprocessing, takes around 1 minute leading to significant time saving and reduced effort for the experimenter. Our ML algorithm is naturally blind to treatment conditions which eliminates the researcher’s confirmation bias. Multiple factors can influence human performance. For instance, with training over time, performance can progressively improve during the scoring phase. However, it can also worsen due to tiredness and distraction when scoring multiple videos consecutively. An ML-based scoring system is immune to these biases, ensuring consistent and standardized analysis.

Another source of bias is the interpretation of animal behaviours, which can vary among researchers leading to differences in classification and quantification. Our ML method provides an unambiguous and standardized tool for the identification of behaviours in the FST.

Compared to the other automatic systems currently available, which require feature extraction, the present 3D RCNN algorithm works directly with video pixels representing a significant advantage. Selecting specific features (such as the degree of mobility, or position coordinates of animal parts) is an arbitrary choice that may cause an oversimplified description of the rodent’s behaviour potentially leading to a loss of valuable information for its identification and labeling.

On the other hand, a potential disadvantage of our model is that it may not perform well in significantly different experimental environments, such as varying light conditions, camera position, or differences in the animal’s size relative to the cylinder. To mitigate these concerns here, we provide a detailed description of the environmental and technical parameters adopted to train and validate our ML model. In addition, it should also be considered that these limitations can be easily overcome by adopting a transfer-learning strategy and retraining the model on new datasets. Indeed, such approaches have proven successful in many computer vision applications (see, e.g., [47]), enabling researchers to address data sparsity for specific tasks by retraining neural networks that were previously trained on more general tasks for which abundant datasets were available.

Another possible strategy is to quickly build a new dataset by manually adjusting the labels predicted by the ML algorithm, following a human-in-the-middle paradigm. Retraining in this way can help refine the algorithm to better suit the specific needs of different laboratories.

In conclusion, we demonstrate that the proposed machine learning model coupled with the specific elaboration of video recordings can effectively discriminate the behavioural effects of different antidepressant drugs. This approach mitigates several biases typically associated with human labeling in recognizing rodents’ behaviours in the forced swim test. Given the structure and adaptability of our ML model, in addition to the FST, it could be applied to the analysis of various behavioural tests in laboratory animals.

## Materials and Methods Animals

A total of 122 male and female Wistar rats were used for the present study. All subjects were bred in-house at the animal facility of the University of Camerino, Italy. Rats were housed four per cage according to their sex and kept under a reversed 12:12h light/dark cycle (lights off at 7 AM) in a temperature (20-22° C) and humidity (45-50%) controlled room. Food (4RF18, Mucedola, Settimo Milanese, Italy) and tap water were provided *ad libitum*.

To train the ML algorithm, 78 male and female rats (n = 39/sex) were used. The first (n = 24, 8/group) and the second experiment (n = 20, 5/group) were conducted in male Wistar rats only.

Before starting the experimental procedures, animals were handled 5 min a day for 5 days by the same operators who performed the experiments. Experiments were conducted during the dark phase of the light/dark cycle. All procedures were approved by the local ethical committee of the University of Camerino and the Italian Ministry of Health (prot.1D580.19). Experiments were conducted in adherence with the *European Community Council Directive for Care and Use of Laboratory Animals and the National Institutes of Health Guide for the Care and Use of Laboratory Animals*.

Experiments were carried out in accordance with ARRIVE 2.0 guidelines.

### Forced Swim Test

The FST experiments were carried out following a previously described and validated procedure [14, 42]. Briefly, swimming sessions were conducted by gently placing rats individually in a transparent plexiglass cylinder (height 50 cm, diameter 28 cm) containing 30-cm water at 23-25° C.

For the drug experiments, two sessions were conducted: an initial 15-min pretest followed 24 hours later by a 5-min test. Drug treatments were administered during the period between these two sessions. Following each swimming session, rats were removed from the cylinder, dried with paper towels, and placed in front of a source of heat for 20 min and then returned to their home cages. Test sessions were videotaped through a camera placed in front of the cylinders (camera resolution: 704 × 576, FPS: 25 Hz). The time spent in immobility, climbing, and swimming was measured.

### Drugs

Fluoxetine hydrochloride (Sigma, St. Louis, MO) and desipramine hydrochloride (Sigma, St. Louis, MO) were dissolved in distilled water and administered subcutaneously (s.c.) at the doses of 20 mg/kg in a volume of 4 ml/kg.

Amitriptyline (E.G. S.p.A, Milan, Italy), and Paroxetine (Angelini S.p.A., Rome, Italy) were diluted in distilled water and administered s.c. at the doses of 20 mg/kg in a volume of 4 ml/kg. Venlafaxine (Italfarmaco S.p.A, Milan, Italy) was prepared as above (diluted in distilled water, 20 mg/kg) and administered s.c. in a volume of 2 ml/kg.

All drugs were administered 23.5, 5, and 1 h prior to the 5-min swimming test [48].

Drug doses were chosen based on published data [49, 14, 40, 41, 42]. To habituate rats to the drug injection procedure they received vehicle injections for three consecutive days before starting the experiments.

### Experimental procedures

#### Dataset preprocessing

To provide the network with all spatiotemporal information necessary for behaviour recognition, as inputs we used Time x Length x Height (*TxLxH*) 3D spatiotemporal tensors, where *T* is the temporal dimension, and *L* and *H* are the spatial dimensions of the resized frame.

The training dataset was composed of 78 videos for a total of 503 minutes, and it was independently labeled by two experienced researchers. Any inconsistency between the two annotators was re-evaluated together to build a univocal dataset.

To label the dataset, the videos were analyzed by identifying the main behaviour in short temporal segments of the video, each lasting 3 seconds. This is a variant of the 5-second scoring method proposed by Slattery and Cryan in 2012 [48].

The length of 3 seconds was chosen to obtain the best balance between an appropriate and clear recognition of the behaviour by the human annotators and dataset dimension that was purposely kept as small as possible to extract more instances from each video collected and limit the computational resources involved.

In this way, 10,062 3D tensors were extracted from the 78 videos.

Swimming, climbing, and immobility were the three main behaviours analyzed. Since diving occurs rarely, it was labeled as swimming [48].

The dataset was preprocessed through a phyton script written using OpenCV module [45, 44, 43] as described below. The area of the cylinder was manually selected, and the resulting Region of Interest (ROI) was wrapped into a rectangle with a standardized dimension *LxH*. To enhance the visibility of the rat every frame was subtracted from the first frame of the video showing no rat. The sequence of those images organized in 3-second blocks was concatenated to build a 3D tensor of size 75×64×128 (*TxLxH*), where 75 is the number of concatenated frames, and 64×128 is the dimensions of a single-wrapped frame. Finally, the pixels of the 3D spatiotemporal tensor were normalized. The whole process is shown in Fig. 8A. The occurrence of each behaviour in the dataset is shown in Fig. 8B.

**Figure 8.**
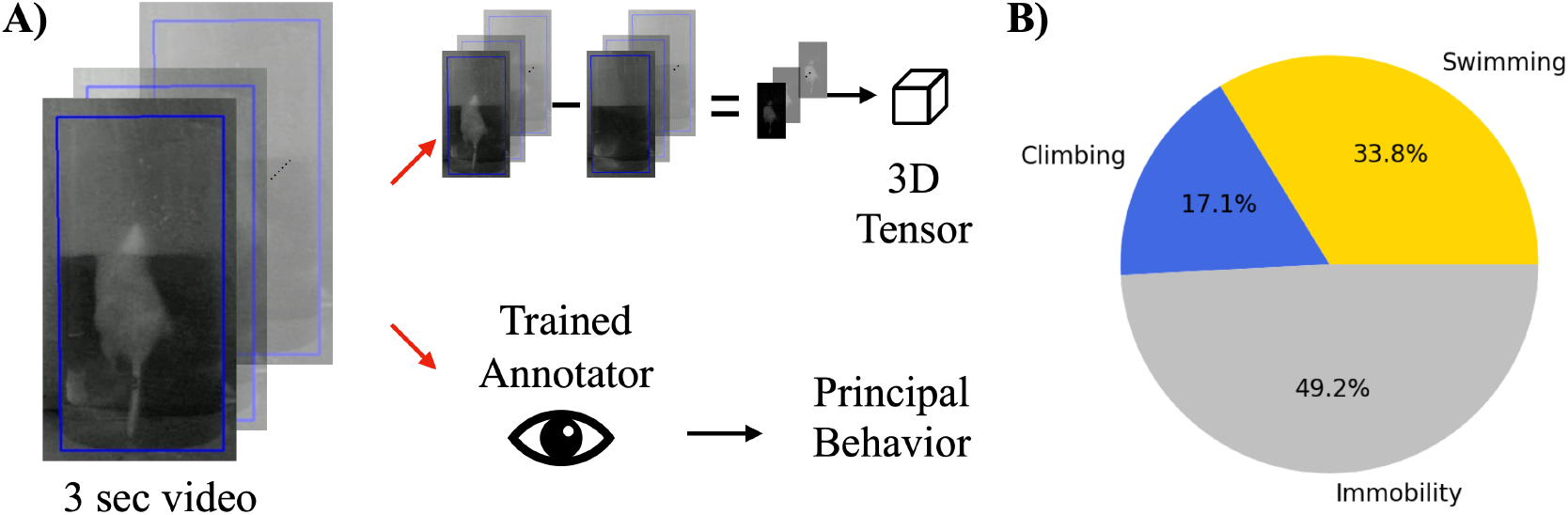
Dataset construction. To build the dataset the video is split into 3-second sub-videos and each sequence of frames composes a single data; every frame is subtracted from the first frame of the video containing no rodent; then, the ROI (blue line) of the cylinder is wrapped into a rectangle of standard dimension (LxH). Finally, the resulting 3D-tensor of dimension TxLxH is normalized. The related portion of the video was manually evaluated by two annotators to recognize immobility, swimming, and climbing behaviours (**A)**. The final dataset is composed of the three interest behaviours (**B**).

The dataset was then randomly split into 90% train and 10% validation sets. The train dataset was used for the optimization of the ML parameters during the training phase, while the validation dataset was used to estimate the performance of the model on new data.

#### Model architecture and training

The model architecture was based on a series of 3D convolutional layers and residual connections, coded using the Keras library [44] with tensorflow backend [45]. The convolutional part was connected to a series of dense layers through a global pooling operation. The role of the dense part was to operate high-level processing on the features extracted by the convolutional layers. Finally, a three-neuron layer with softmax activation was used for the recognition of the behavioural classes. Since the behaviour recognition is invariant under the mirroring of the images around the vertical axis, the training dataset was batch-augmented by making use of this transformation [50].

The model was trained for 120 epochs exploiting computational acceleration provided by a state-of-the-art graphic processing unit, namely, a GPU RTX A6000. To prevent the model from overfitting on the train dataset, early stopping was used. The model showing the highest accuracy on the validation set was selected and then used for pharmacological validation experiments.

The library versions used are reported in Table 2.

**Table 2.** The version of the main libraries used. The main Python libraries used to develop the code in the following rows.

| Python Library | Version |
| --- | --- |
| Tensorflow | 2.11.0 |
| Keras | 2.11.0 |
| Numpy | 1.24.2 |
| CUDA | 12.2 |
| Python | 3.9.9 |
| OpenCV | 4.5.2 |

### Comparison of the ML algorithm with the human manual scoring: Evaluation of the forced swimming effects of FLX and DMI

To evaluate the accuracy and reliability of the ML algorithm trained, a pharmacological experiment was performed to test the forced swimming response to established antidepressant drugs. Two groups of rats subjected to a 15-min pretest received FLX (20 mg/kg, 4 ml/kg) or DMI (20 mg/kg, 4 ml/kg) 24 h, 5 h, and 1 h prior to the start of the given 5-min test. A third group of animals was treated with saline and served as a control. Videos were analyzed both by one of the two experienced annotators and the trained ML algorithm scoring system.

As previously described, the videos were split into 3 seconds sub-video and preprocessed. The resulting 3D tensor was analyzed by the trained model (Fig. 9).

**Figure 9.**
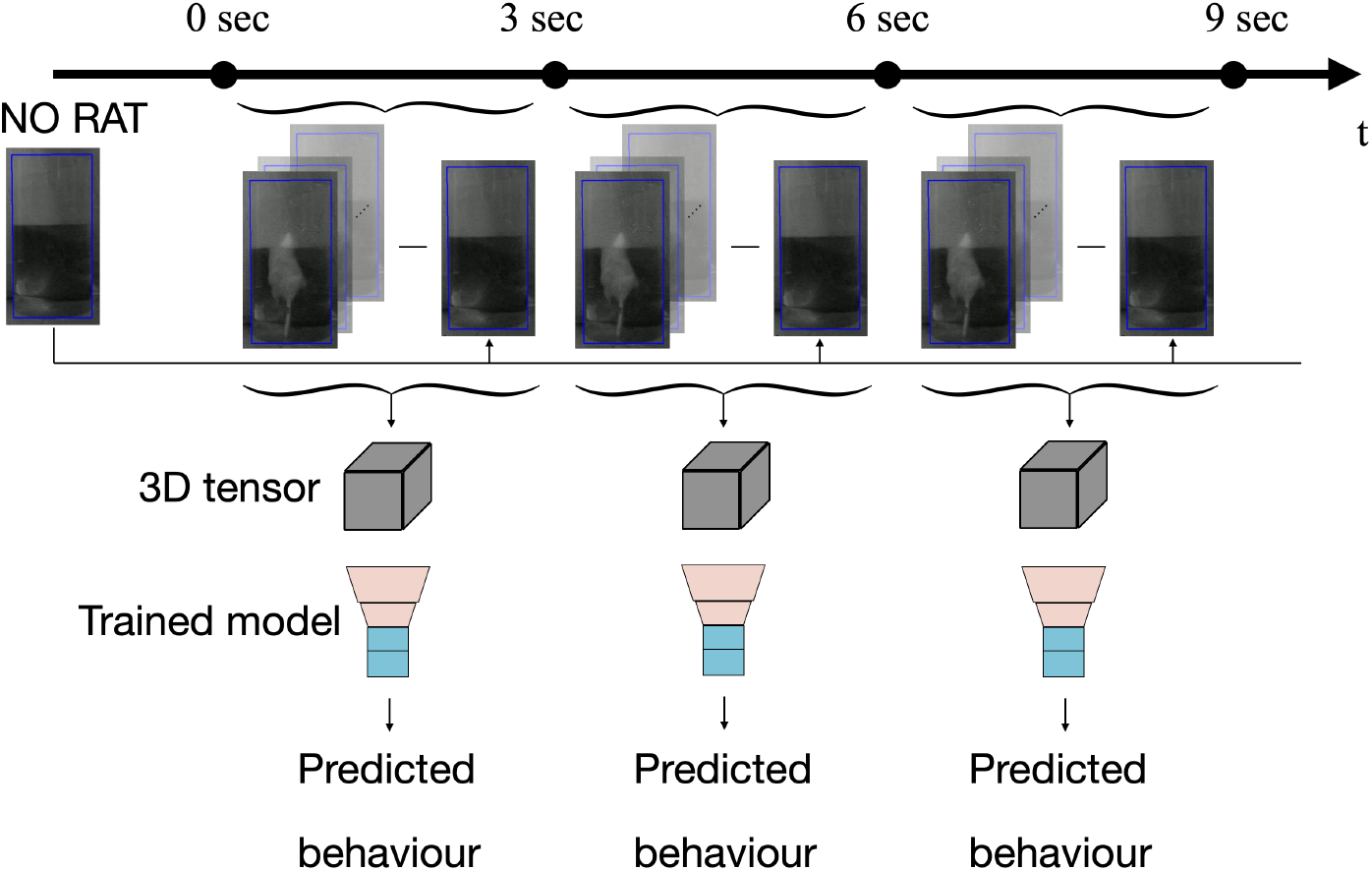
Pipeline of ML annotation applied to a video. The video is split into 3-second sub-videos, and they are preprocessed as already described in Fig. 8: every frame of the sequence is subtracted from the first frame of the original video containing no rat; then, the ROI (blue line) of the cylinder is wrapped into a rectangle of standard dimension (LxH). Finally, the resulting 3D-tensor of dimension TxLxH is normalized. The resulting 3D tensor is evaluated by the model already trained.

### ML algorithm evaluation of the effects of AMI, PRX, and VLX in the FST

A second pharmacological experiment was carried out to assess the effectiveness of our ML algorithm in distinguishing the subtle differences in the swimming response elicited by different classes of antidepressants. For this purpose, employing the same protocol described above, AMI (20 mg/kg, 4 ml/kg), PRX (20 mg/kg, 4 ml/kg), and VLX (20 mg/kg, 2 ml/kg) were used.

### Statistical analysis

Behavioural data were analyzed by ANOVA using GraphPad Prism version 9.5.1 (GraphPad Software, San Diego, California, USA).

In the first experiment, the comparison between annotator and ML scoring methods was evaluated for each behaviour via two-way ANOVA with scoring methods and treatment as between factors. In the second experiment, when the only ML scoring method was applied, the one-way ANOVA was used with drug as between-subjects factor.

ANOVA was followed by the Dunnett’s post-hoc test when appropriate, and statistical significance was conventionally set at p < 0.05.

To evaluate the correlation of the two scoring methods the Pearson’s correlation was used.

## Data availability

The datasets generated and analyzed and the code are available in the GitHub repository [51] and the Zenodo Repository of Ref. [52].

## Acknowledgments

We thank Sofia De Cillis for her help in carrying out the FST experiments, Rina Righi and Agostino Marchi for animal care. The computer used to train the ML model was provided by the Physics Division of the University of Camerino. We acknowledge contributions from the following projects: PRIN-PNRR 2022 MUR project “UEFA”-P2022NMBAJ, PRIN-PNRR2022 - P202274WPN (to RC), PRIN-2022 20227HRFPJ (to RC), PRIN-2022 2022H77XB7, NGEU+MUR Award ID: project MNESYS (PE0000006)-Project AMSUD 2024 “Animal Models to Study Substance Use Disorders and Cooccurring Psychiatric Conditions”.

## References

[1] American Psychiatric Association. Diagnostic and Statistical Manual of Mental Disorder, 5-TR ed. (2022).

[2] W. H. Organization. The global burden of disease: 2004 update https://iris.who.int/bitstream/handle/10665/43942/9789241563710_eng.pdf?sequence=1 (2004).

[3] W. H. Organization. Depressive disorder (depression) https://www.who.int/news-room/fact-sheets/detail/depression (2023).

[4] Fornaro, M. et al. The emergence of loss of efficacy during antidepressant drug treatment for major depressive disorder: an integrative review of evidence, mechanisms, and clinical implications. Pharmacol Res. 139, 494–502 (2019).

[5] Licinio, J. & Wong, M. L. Depression, antidepressants and suicidality: a critical appraisal. Nat Rev Drug Discov. 4, 165–171 (2005).

[6] Penn, E. & Tracy, D. K. The drugs don’t work? antidepressants and the current and future pharmacological management of depression. Ther Adv Psychopharmacol. 2, 179–188 (2012).

[7] Commons, K. G., Cholanians, A. B., Babb, J. A. & Ehlinger, D. G. The rodent forced swim test measures stress-coping strategy, not depression-like behavior. ACS Chem Neurosci. 8, 955–960 (2017).

[8] Porsolt, R. D. Behavioral despair in Antidepressant, Neurochemical, Behavioral and Clinical Perspectives (eds. Enna, S. J., Malick, J.B. & Richelson, R.) 121–139 (Raven Press, 1981).

[9] Borsini F. & Meli, A. Is the forced swimming test a suitable model for revealing antidepressant activity? Psychopharmacology (Berl). 94, 147–160 (1988).

[10] Lucki, I. The forced swimming test as a model for core and component behavioral effects of antidepressant drugs. Behav Pharmacol. 8, 523–532 (1997).

[11] Kara, N. Z., Stukalin, Y. & Einat, H. Revisiting the validity of the mouse forced swim test: systematic review and meta-analysis of the effects of prototypic antidepressants. Neurosci Biobehav Rev. 84, 1–11 (2018).

[12] Petit-Demouliere, B., Chenu, F. & Bourin, M. Forced swimming test in mice: a review of antidepressant activity. Psychopharmacology (Berl). 177, 245–255 (2005).

[13] Cryan, J. F., Valentino, R. J., & Lucki, I. Assessing substrates underlying the behavioral effects of antidepressants using the modified rat forced swimming test. Neurosci Biobehav Rev. 28, 547–569 (2005).

[14] Detke, M. J., Rickels, M. & Lucki, I. Active behaviors in the rat forced swimming test differentially produced by serotonergic and noradrenergic antidepressants. Psychopharmacology (Berl). 121, 66–72 (1995).

[15] Detke, M. J. & Lucki, I. Detection of serotonergic and noradrenergic antidepressants in the rat forced swimming test: the effects of water depth. Behav Brain Res. 73, 43–46 (1996).

[16] Hollis, F. & Kabbaj, M. Social defeat as an animal model for depression. ILAR J. 55, 221–232 (2014).

[17] Katz, R. J., Roth, K. A., & Carroll, B. J. Acute and chronic stress effects on open field activity in the rat: implications for a model of depression. Neurosci Biobehav Rev. 5, 247–251 (1981).

[18] Rygula, R. et al. Anhedonia and motivational deficits in rats: impact of chronic social stress. Behav Brain Res. 162, 127–134 (2005).

[19] Sahin, Z. et al. Investigation of effects of two chronic stress protocols on depression-like behaviors and brain mineral levels in female rats: an evaluation of 7-day immobilization stress. Biol Trace Elem Res. 199, 660–667 (2021).

[20] Venzala, E., García-García, A. L., Elizalde, N. & Tordera, R. M. Social vs. environmental stress models of depression from a behavioural and neurochemical approach. Eur Neuropsychopharmacol. 23, 697–708 (2013).

[21] Armario, A. The forced swim test: Historical, conceptual and methodological considerations and its relationship with individual behavioral traits. Neurosci Biobehav Rev. 128, 74–86 (2021).

[22] de Kloet, E. R. & Molendijk, M. L. Coping with the forced swim stressor: towards understanding an adaptive mechanism. Neural Plast. 2016, 6503162; 10.1155/2016/6503162 (2016).

[23] Molendijk, M. L. & de Kloet, E. R. Forced swim stressor: trends in usage and mechanistic consideration. Eur J Neurosci. 55, 2813–2831 (2022).

[24] Bohnslav, J. P. et al. DeepEthogram, a machine learning pipeline for supervised behavior classification from raw pixels. Elife. 10, e63377; 10.7554/eLife.63377 (2021).

[25] Gulinello, M. et al. Rigor and reproducibility in rodent behavioral research. Neurobiol Learn Mem. 165, 106780; 10.1016/j.nlm.2018.01.001 (2019).

[26] Segalin, C. et al. The mouse action recognition system (MARS) software pipeline for automated analysis of social behaviors in mice. Elife. 10, e63720; 10.7554/eLife.63720 (2021).

[27] Crowley, J. J., Jones, M. D., O’Leary, O. F. & Lucki, I. Automated tests for measuring the effects of antidepressants in mice. Pharmacol Biochem Behav. 78, 269–274 (2004).

[28] Fitzgerald, P. J., Yen, J. Y. & Watson, B. O. Stress-sensitive antidepressant-like effects of ketamine in the mouse forced swim test. PLoS One. 14, e0215554; 10.1371/journal.pone.0215554 (2019).

[29] Hédou, G., Pryce, C., Di Iorio, L., Heidbreder, C. A. & Feldon, J. An automated analysis of rat behavior in the forced swim test. Pharmacol Biochem Behav. 70, 65–76 (2001).

[30] Juszczak, G. R. et al. The usage of video analysis system for detection of immobility in the tail suspension test in mice. Pharmacol Biochem Behav. 85, 332–338 (2006).

[31] Nandi, A., Virmani, G., Barve, A. & Marathe, S. DBscorer: An open-source software for automated accurate analysis of rodent behavior in forced swim test and tail suspension test. eNeuro. 8, ENEURO.0305-21.2021; 10.1523/ENEURO.0305-21.2021 (2021).

[32] Sturman, O. et al. Deep learning-based behavioral analysis reaches human accuracy and is capable of outperforming commercial solutions. Neuropsychopharmacology. 45, 1942–1952 (2020).

[33] Yuman, N. et al. High-speed video analysis of laboratory rats behaviors in forced swim test. 2008 IEEE International Conference on Automation Science and Engineering. 206–211 (2008).

[34] von Ziegler, L., Sturman, O. & Bohacek, J. Big behavior: challenges and opportunities in a new era of deep behavior profiling. Neuropsychopharmacology. 46, 33–44 (2021).

[35] Perez, M. & Toler-Franklin, C. CNN-based action recognition and pose estimation for classifying animal behavior from videos: a survey. Preprint at https://arxiv.org/abs/2301.06187 (2023).

[36] Tran, D., Bourdev, L., Fergus, R., Torresani, L. & Paluri, M. Learning spatiotemporal features with 3D convolutional networks. Preprint at https://arxiv.org/abs/1412.0767 (2015).

[37] He, K., Zhang, X., Ren, S. & Sun, J. Deep residual learning for image recognition. Preprint at https://arxiv.org/abs/1512.03385 (2015).

[38] Detke, M. J., Johnson, J. & Lucki, I. Acute and chronic antidepressant drug treatment in the rat forced swimming test model of depression. Exp Clin Psychopharmacol. 5, 107–112 (1997).

[39] Rex, A., Schickert, R. & Fink, H. Antidepressant-like effect of nicotinamide adenine dinucleotide in the forced swim test in rats. Pharmacol Biochem Behav. 77, 303–307 (2004).

[40] Enríquez-Castillo, A. et al. Differential effects of caffeine on the antidepressant-like effect of amitriptyline in female rat subpopulations with low and high immobility in the forced swimming test. Physiol Behav. 94, 501–509 (2008).

[41] Flores-Serrano, A. G. et al. Clinical doses of citalopram or reboxetine differentially modulate passive and active behaviors of female Wistar rats with high or low immobility time in the forced swimming test. Pharmacol Biochem Behav. 110, 89–97 (2013).

[42] Rénéric, J. P. & Lucki, I. Antidepressant behavioral effects by dual inhibition of monoamine reuptake in the rat forced swimming test. Psychopharmacology (Berl). 136, 190–197 (1998).

[43] OpenCV. Open Source Computer Vision Library https://opencv.org/ (2015).

[44] Gulli, A. & Pal, S. Deep learning with Keras: Implementing Deep Learning Models and Neural Networks with the Power of Python (Packt Publishing Ltd, 2017).

[45] Abadi, M. et al. Large-scale machine learning on heterogeneous distributed systems. Preprint at https://arxiv.org/abs/1603.04467 (2015).

[46] Kingma, D. P. & Ba, J. Adam: a method for stochastic optimization. Preprint at https://arxiv.org/abs/1412.6980v9 (2017).

[47] Hussain, M., Bird, J. J. & Faria, D. R. A study on CNN transfer learning for image classification in Advances in Intelligent Systems and Computing (eds. Lotfi, A., Bouchachia, H., Gegov, A., Langensiepen, C., McGinnity, M.) 840, 191–202 (Springer, Cham, 2019).

[48] Slattery, D. A. & Cryan, J. F. Using the rat forced swim test to assess antidepressant-like activity in rodents. Nat Protoc. 7, 1009–1014 (2012).

[49] Cryan, J. F. & Lucki, I. Antidepressant-like behavioral effects mediated by 5-Hydroxytryptamine(2C) receptors. J Pharmacol Exp Ther. 295, 1120–1126 (2000).

[50] Hoffer, E., et al. Augment your batch: better training with larger batches. Preprint at https://arxiv.org/abs/1901.09335 (2019).

[51] Della Valle, A., De Carlo, S., De Cillis, S., Sonsini, G., Pilati, S., Perali, A., Ubaldi, M. & Ciccocioppo, R. 3D residual convolutional neural network for forced swim test dataset. GitHub https://github.com/adso42/FST_3DRCNN (2025).

[52] Della Valle, A., De Carlo, S., De Cillis, S., Sonsini, G., Pilati, S., Perali, A., Ubaldi, M. & Ciccocioppo, R. Data and code for “Machine-learning based model for behavioral analysis in rodents: application to the forced swim test” dataset. Zenodo https://zenodo.org/records/14638257 (2025).

